# Development of a Reproducible Porcine Model of Infected Burn Wounds

**DOI:** 10.1101/2021.08.16.456568

**Authors:** Sayf Al-deen Said, Samreen Jatana, András K. Ponti, Erin E. Johnson, Kimberly A. Such, Megan T. Zangara, Maria Madajka, Francis Papay, Christine McDonald

## Abstract

Severe burns are traumatic and physically debilitating injuries with a high rate of mortality. Bacterial infections often complicate burn injuries, which presents unique challenges for wound management and improved patient outcomes. Currently, pigs are used as the gold standard of pre-clinical models to study infected skin wounds due to the similarity between porcine and human skin in terms of structure and immunological response. However, utilizing this large animal model for wound infection studies can be technically challenging and create issues with data reproducibility. We present a detailed protocol for a porcine model of infected burn wounds based on our experience in creating and evaluating partial thickness burn wounds infected with *Staphylococcus aureus* on six pigs. Wound healing kinetics and bacterial clearance were measured over a period of 27 days in this model. Enumerated are steps to achieve standardized wound creation, bacterial inoculation, and dressing techniques. Systematic evaluation of wound healing and bacterial colonization of the wound bed is also described. Finally, advice on animal housing considerations, efficient bacterial plating procedures, and overcoming common technical challenges is provided. This protocol aims to provide investigators with a step-by-step guide to execute a technically challenging porcine wound infection model in a reproducible manner. Accordingly, this would allow for the design and evaluation of more effective burn infection therapies leading to better strategies for patient care.

**Graphical Abstract:** 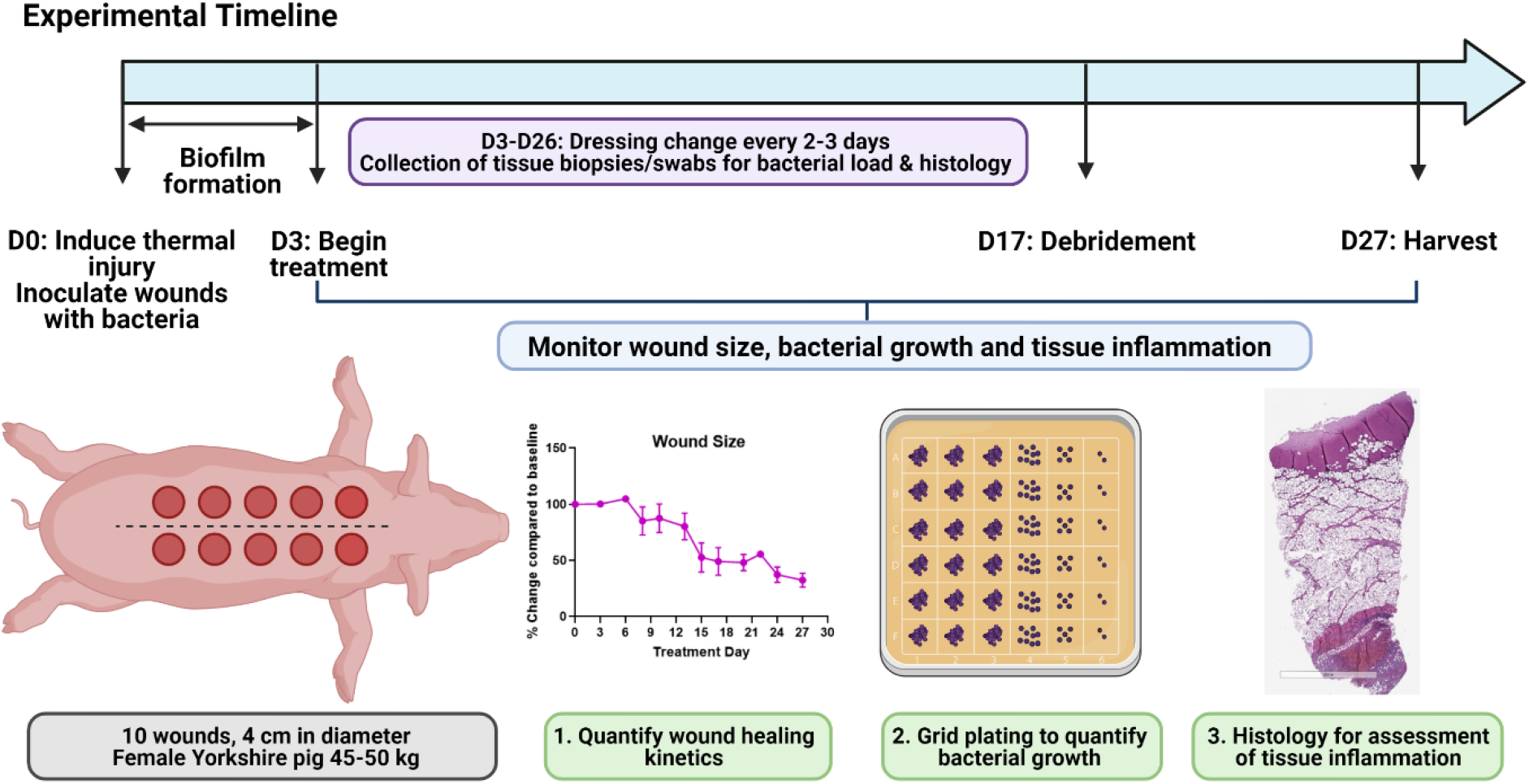

## Background

Burn injuries constitute a major clinical and economic challenge for healthcare systems worldwide. Approximately 11 million burn injuries of varying severity occur annually worldwide leading to 180,000 total fatalities as estimated by the World Health Organization [1]. Severe burn wounds cause both physical and psychological trauma resulting in decreased quality of life. Burn injuries interfere with the normal wound healing process and alter the immune responses by disrupting the anatomical and immunological barriers to infection [2, 3]. Wound sites are frequently infected with drug resistant bacterial strains, like methicillin-resistant *Staphylococcus aureus* (MRSA), resulting in high morbidity and mortality [3-5]. Therefore, it is imperative to develop relevant and reproducible pre-clinical models that allow investigators to study complications, such as bacterial infections, occurring as a result of burn injuries and test the development of novel therapeutic approaches that optimize patient outcomes improving the overall standard of care.

The porcine model represents the most relevant animal model to study wound infection due to the similarities between porcine and human skin in structure, immunological response, and healing patterns [6-10]. The stratum corneum thickness, vascularization, keratin types, and extracellular matrix composition of porcine and human skin are similar, presenting similar elastic components that are important in wound healing. However, this type of large animal model presents with unique technical nuances and challenges that can significantly impact the study outcomes and data reproducibility. Unlike small animal studies, the financial costs and personnel effort associated with large animal studies precludes the ability to rapidly optimize protocols or easily increase the number of animals utilized to obtain statistical power. In addition, infection control measures in a large animal setting are very different from small animal models and require additional considerations to ensure personnel safety and limit contamination with infectious agents.

This protocol provides detailed information to implement a porcine model of infected burn wounds and evaluate wound healing and infection parameters. The protocol is based on data collected from performing second-degree, partial thickness, circular burn wounds (n=10 wounds/pig) on the back of female Yorkshire pigs (40-50kg, n=6 total animals) followed by inoculation with *Staphylococcus aureus*. We utilized this particular strain in this study due to the high prevalence of *S. aureus* associated infections in skin wounds. Wounds were monitored for healing kinetics and bacterial colonization over a period of 27 days (**Fig. 1**). Enumerated are steps to achieve a standardized porcine wound infection model that can be utilized by investigators to design and test effective therapeutics that treat bacterial infections. This protocol serves as an important resource for pre-clinical testing in a large animal setting prior to testing a drug or a therapeutic in patients suffering from similar injuries and infections in clinical trials.

**Figure 1:**
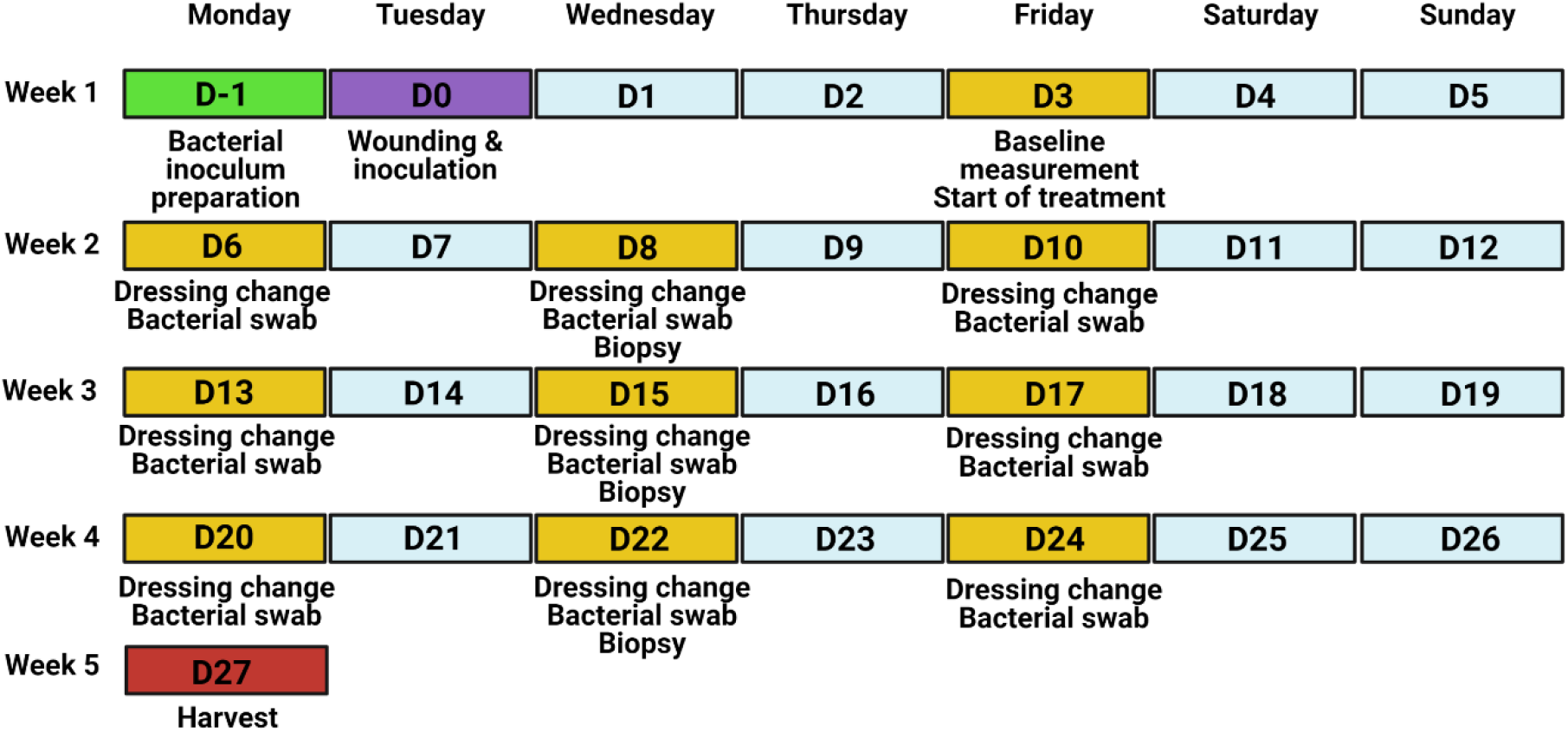
Experimental timeline of the porcine wound infection model.

## MATERIALS

### Reagents & Supplies

- *Staphylococcus aureus* RN4220 [11] or *S. aureus* (American Type Culture Collection, Cat. # ATCC® 25923)
- Tryptic soy broth (TSB) (Fisher Scientific, Cat. # R455052)
- Luria Bertrani (LB) agar (Fisher Bioreagents, Cat. # BP1425-500)
- BBL mannitol salt agar (MSA) (BD Biosciences, Cat. # 211407)
- 100 mm square polystyrene Petri dishes (Fisher Scientific, Cat. # FB0875711A)
- 14 mL polystyrene round bottom tubes (Corning, Cat. # 352057)
- 96 well flat bottom plates (USA Scientific, Cat. # 5665-5161)
- Cotton tipped applicators (Medline, Cat. # MDS202000)
- Integra™ Miltex™ 3 mm biopsy punches with plunger system (Fisher Scientific, Cat. # 12-460-407)
- HistoChoice® MB tissue fixative (Sigma-Aldrich, Cat. # H2904)
- Betadine® solution, 7.5% povidone-iodine (Betadine, Cat. # BSO16P)
- Triple pack alcohol swab sticks (Medline, Cat. # MDS093810)
- Avant gauze non-woven gauze sponges 3”x3” (Medline, Cat. # NON21334)
- Telfa™ Adhesive Island dressing, 4”×4” (Medtronic, Cat. # COVH7550)
- Tincture of benzoin (3M, Cat. # MC1544)
- Steri-Drape™ (3M, Cat. # M1050)
- Elastikon® elastic tape (Johnson & Johnson, Cat. # JJ-005175)
- Weck Visistat skin stapler 35W (Teleflex, Cat. # W528235)
- Weckstat skin staple remover (Teleflex, Cat. # W525980)
- Stainless steel scalpel #10 (Medline, Cat. # MDS15210)
- Blank stencil sheets, 4 mil mylar 12”x12” (Stencil Ease, SKU AMY041212-012-M004)
- 50 mL centrifuge tubes (Corning, Cat. # 430829)
- Natural cork underlayment Roll (QEP, Cat. # 72003Q)
- Push pins (Staples, Cat. # 10552)

### Equipment

- GENESYS 30 visible spectrophotometer with test-tube holder (Fisher Scientific, Cat. # 14385340 & 14380443)
- Labnet mini-incubator (Laboratory Product Sales, Cat. # I-5311-DS)
- Custom Burn Instrument: An in house manufactured device with temperature controlled burn head, pressure sensor, and timer (plans available upon request; **Supplementary Fig. 1**). The main components include a thermal device (Wallbrand™ Electric Branding Tools, Hand-held (Wall Lenk Corp., Cat. # WB200), a heat sensor (SOLO Single-Loop Temperature Controller 1/16 DIN; Solo Inc., Cat. # SL4848-RR), a pressure sensor (ProSense Series Panel Meter; Pro·Sense®, Cat. # DPM3-AT-2R-H), and a digital timer (AutomationDirect, Cat. # CTT-1C-A120)
- Ekare inSight 3D (eKare Inc., Fairfax, VA)
- Iris Surgical Scissors 4½” Straight Stainless (Teleflex, Cat. # 144300)
- Adson Tissue Forceps 4¾” (Teleflex, Cat. # 181223)
- Metzenbaum Dissecting Scissors (Teleflex, Cat. # 464715)

## PROCEDURE

1. **Bacterial inoculum preparation**
  1.1 Prepare a fresh bacterial stock plate on selective agar and incubate overnight at 37°C.
  1.2 Inoculate 4 mL TSB broth with a single *S. aureus* colony and incubate overnight at 37°C, shaking at 220 rpm.
  1.3 Measure the optical density at 600 nm (OD_600_) of the overnight culture using a spectrophotometer.
  1.4 Inoculate fresh TSB broth with the overnight culture at a 1:10 dilution (0.4 mL overnight culture into 3.6 mL TSB broth) and incubate at 37°C for 1 hour at 220 rpm. Measure OD_600_ of the culture at the start of incubation (0 h).
  1.5 After 1 hour, measure the OD_600_ of the secondary culture to confirm log phase growth and calculate the CFU/µL (see NOTE (iii) for more detail).
  1.6 Prepare the inoculum to be added to the wounds in sterile PBS. Calculate the volume of secondary culture required to inoculate each wound with 10 ^6^ CFU and pellet the bacteria by centrifugation at 1,500 rpm for 15 minutes. Carefully remove the supernatant and re-suspend the bacterial pellet in sterile PBS at the desired concentration (e.g. a concentration of 3.3×10^6^ CFU/mL to deliver 10^6^ CFU in a volume of 0.3 mL to a 4 cm diameter wound).
  1.7 Plate the inoculum in duplicate on selective agar plates for confirmation of CFU applied to the wounds. Refer to section 6 for the plating procedure. **TIPS:** (i) Set up replicate overnight cultures (e.g., 2-3 cultures) to ensure sufficient bacteria culture for use the following day. (ii) Always prepare excess inoculum (e.g., 4 times the required volume) to facilitate efficient pelleting and ease of administration (Step. 1.6). **HINT :** An example of inoculum calculation
    - If OD_600_ of secondary culture = 0.452 and correlates with 1.94×10^5^ CFU/µL based on the growth curve, then 10^6^ CFU = 5.15 µL culture/wound
    - If creating 10 wounds total, 10 wounds x 5.15 µL/wound = 51.5 μL total
    - Will need to generate 4 times the calculated inoculum, 51.5 μL x 4 = 206 µL
    - Centrifuge 206 µL of secondary culture at 1,500 rpm. Resuspend pellet in 12 mL sterile PBS (0.3 mL/wound x 10 wounds x 4 = 12 mL).
    **NOTE**: (i) The specific bacterial strain used to inoculate the burn wounds may vary depending on the research question and institutional restrictions. Standard microbiological procedures are used for bacterial culture and BSL2 safety procedures are employed for the kanamycin-resistant *S. aureus* strain used in this protocol. Consult with your local Institutional Biosafety Committee for more information about recommended practices and procedures for handling BSL2 organisms. (ii) Selective agar refers to agar plates with antibiotics to select for the specific bacterial strain (e.g. LB with kanamycin). Selective and differential agar that permits the growth and identification of specific types of bacteria (e.g. MSA for *Staphylococcus* strains, fermentation of mannitol by *S. aureus* turns it yellow). (iii) Prior to the start of an animal experiment, bacterial growth rate, correlation between culture density and CFU and appropriate selective and/or differential media will need to be determined to aid in the calculations utilized for animal inoculation and evaluation of bacterial colonization. (iv) The optimal inoculation of 4 cm wounds is 10^6^ CFU in 0.3 mL volume. The bacterial amount and volume should be scaled proportionally to fit different size burn wounds.
2. **Administration of anesthesia**
  2.1 Sedate the animal with an intramuscular injection of 4.4-6.6 mg/kg of Telazol.
  2.2 Intubate the pig and place on 1-3% isoflurane inhalation to maintain sedation.
  2.3 Inject 0.01-0.05 mg/kg buprenorphine intramuscularly to control pain during the wounding procedure.
  2.4 Place a fentanyl transdermal patch (75 μg) on the upper and posterior part of the neck and secure with a small piece of Elastikon® and skin staples (do not staple through patch). Replace every 72 hours.
  **NOTE**: (i) To minimize the risk of aspiration during anesthesia, fast the pig overnight with access to only water. (ii) Although transdermal fentanyl is the primary means of pain control during the experiment, its analgesic effect does not start until 12 hour post-administration. Buprenorphine is administered to control pain during this 12 hour lag time. (iii) The interval between dressing changes (2-3 days) will determine the dose and duration of the fentanyl patch, as it is easier to place and secure during sedation. (iv) If NSAIDs, such as meloxicam or carprofen, are not expected to interfere with experimental outcomes they should be considered for use after the initial burn procedure in consultation with veterinary staff.
3. **Burn wound creation and inoculation**
  3.1 Under anesthesia, shave the back and sides of the pig with an electric razor. Clean off the back with a moist cloth to remove hair before transferring the pig to the operating room.
  3.2 Position the pig prone on a procedure table covered with disposable, absorbent pads and mark the wound positions using the prepared plastic template and skin marker. **CRITICAL STEP:** Pay special attention to the positioning of the pig, as any spine curvature can impact the correct placement of wounds, alter the skin tension of the wounds, and cause variation in the rate of wound healing.
  3.3 Clean and prepare the burn area using sterile gauze soaked in betadine solution working from the spinal area to the sides. Cover the pig with sterile drapes.
  3.4 Create burn wounds according to the layout shown in **Fig. 2** using a controlled pressure and temperature device (**Supplementary Fig. 2**). To achieve a partial thickness burn on a 45-50 kg Yorkshire pig, set burn temperature to 200°C and apply pressure of 11.9 PSI for 60 seconds.
  3.5 Document the baseline wound area by taking photographs of individual wounds with a reference scale.
  3.6 Allow wounds to cool before adding the bacterial inoculum. Using a p1000 pipette and barrier tips, slowly add 0.3 mL bacterial inoculum dropwise over the surface of the wound and using a sterile cotton swab.
  **TIP:** Prior to the experiment, confirm that the temperature, pressure, and time settings create the depth of wound desired with your burn device using cadaver pig skin *ex-vivo*. **NOTE**: (i) The elbow and the stifle joints should be positioned to align the pig close to a natural position. This positioning should be replicated in future dressing changes (Step 3.2). (ii) Commercial electric branding irons can be used instead of the custom device shown here; however, they will not provide information about pressure applied during burning and may result in more variable wound creation. (iii) We found that the eKare inSight 3D system provides quick, accurate, and consistent measurements of wound area. (iv) Mix the inoculum gently but thoroughly, before administering, as the bacteria may settle to the bottom of the tube. This creates a uniform suspension such that consistent amounts of bacteria are administered to each wound (Step 3.6). (v) Position a sterile gauze pad at the lower edge of the wound to prevent any drips during inoculation that would contaminate the skin or work surface (Step 3.6). This is most easily performed with a second person (one to inoculate and the other to hold the gauze).
4. **Wound dressing application**
  4.1 Cover each wound with a 3”x3” sterile, non-stick gauze pad with the gauze fold positioned medially (**Fig. 2C**).
  4.2 Apply tincture of benzoin adhesive solution around the gauze pads (**Fig. 2D**).
  4.3 Cover gauze pad with 4”×4” Telfa™ dressing and secure with four staples. The staples need to oppose each other to maintain tension on the Telfa™ dressing (**Fig. 2E**).
  4.4 Apply three long Elastikon® strips to secure the dressings; one along the spine so that it covers the medial portion of the dressing on both sides and the other two on the side of the torso to cover the lateral side of the dressings (**Fig. 2F**).
  4.5 Staple through Elastikon® strips to further reinforce each Telfa™ bandage (**Fig. 2F**).
  4.6 Apply Steri-Drape™ bandage over the entire back. This layer will minimize contamination, even if the underlying gauze collects excessive discharge.
  4.7 Anchor the Steri-Drape™ through the addition of Elastikon® strips across the back and wrapping around the shoulders and hips (**Fig. 2G**).
  **TIPS**: (i) One tube of tincture of benzoin is sufficient for two wounds and is most adhesive while wet; therefore, best results occur when the Telfa™ dressing is secured shortly after application of tincture of benzoin. (ii) If the Steri-Drape™ extends to cover forelimbs or hind limbs, cut the segment behind the forelimb or in front the hind limbs to allow free movement of the limbs. **NOTE**: (i) Monitoring the animal at least twice a day to assess for early signs of dressing failure is paramount to the success of the experiment. As wound healing progresses, animals may rub against enclosure walls to either try to remove the dressing, or alleviate itching. Addressing such shortfalls promptly by placing patches of Steri-Drape™ or reinforcing with Elastikon® is generally enough to keep the dressing intact until the next dressing change. (ii) To minimize incidence of wound dressing failure, the animal must be housed in a cage free of angular protrusions (e.g. screw heads, automated water systems, etc.) that can catch on the bandage while the animal rubs against the wall.
5. **Dressing changes and sample collection**
  5.1 Sedate the pig with an intramuscular injection of Telazol (4.4-6.6 mg/kg). Intubate and maintain anesthesia by 1-3% isoflurane inhalation.
  5.2 Remove the anchoring Elastikon strips and other layers of the dressing using a surgical staple remover.
  5.3 Carefully clean the skin using sanitary wipes, working from the outer areas in towards the wounds to remove wound exudate, residual adhesive, and other detritus.
  5.4 Carefully clean the area adjacent to the wounds with 70% ethanol swabs to aid in visualization of the wound edge and photograph individual wounds.
  5.5 Collect a sample of bacteria colonizing the wound surface. Wet the tip of a sterile cotton-tipped applicator with the PBS (0.5 mL) in the collection tube. Gently roll the swab over the entire surface of the wound. Return the swab to the collection tube, cap the tube, and place on ice.
  5.6 Using a 3 mm biopsy punch, collect tissue from each wound and place into labeled containers with fixative (e.g. HistoChoice®) for subsequent paraffin embedding and histological analysis.
  5.7 Remove necrotic tissue in the third week of the experiment by lifting the wound eschar with sterile Adson forceps and iris scissors to separate the eschar from the wound bed.
  **NOTE**: (i) Culture analysis to quantify CFU should be performed promptly after sample collection for best results. This can be achieved by incorporating a dedicated team member to retrieve and process the samples immediately after collection. (ii) Tissue biopsy samples should be collected from the same region from the separate wounds (Step 5.6). For example, if the wound is a clock face, at one collection point all samples will be taken at the 12 o’clock position. (iii) Make the debridement day consistent (e.g. D17) between experiments, as this significantly alters wound appearance and the wound surface available for viable bacteria quantification (Step 5.7).
6. **Evaluation of bacterial colonization of wounds**
  6.1 In a biosafety cabinet, prepare a sterile 96 well plate with 80 µL sterile PBS in columns 2-2 using a multichannel pipettor (**Fig. 3A**).
  6.2 Vortex the collection tubes and then transfer 100 µL to the first column of a sterile 96-well plate in duplicate (**Fig. 3A**).
  6.3 Serially dilute the sample 1:5 across the rows of the plate (**Fig. 3A**). **CRITICAL STEP:** Change pipet tips for each dilution (between columns) to prevent bacterial aggregation.
  6.4 Using a multichannel-pipette, transfer 5 µL of each dilution to the surface of a bacterial plate to create a small spot (Fig. 3B). Perform this on duplicate agar plates (e.g., 4 spots total on 2 plates). Start with the lowest dilution (column 12) and work back to the undiluted sample (column 1). **CRITICAL STEP:** Change pipette tips between each column during spot plating to prevent bacterial aggregation.
  6.5 Keep agar plates upright on the benchtop until dry (e.g., the spots are absorbed into the agar), then invert and transfer to a 37°C incubator for growth overnight.
  6.6 Count the colonies in each spot for each dilution. Total the colonies from the duplicate samples on duplicate plates (4 spots/wound).
  6.7 Calculate the CFU/wound using the following equation:

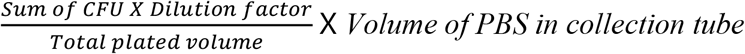
  **TIPS**: (i) Label all plates the night before collection to reduce time between sample procurement and plating. (ii) Remove pipet tips from the 3^rd^ and 6^th^ row of the box to facilitate the transfer and serial dilution of samples using this plate set up (Step 6.3). (iii) Skip rows C and F to allow for spacing between samples on the culture plate (Step 6.1). **HINT** : An example for CFU calculation if you plated a 1:5 dilution series and the bacteria were counted at the 7^th^ dilution for 4 spots plated in duplicate:

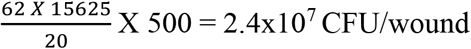

62 is the total colonies from 4 spots 15625 is the dilution factor (7^th^ dilution) 20 is total plated volume (5 μL plated in 4 spots) 500 μL is the volume of PBS in the collection tube **NOTE**: (i) For any new experiment, the dilution might need to be optimized. A 1:10 and 1:5 dilution series at the initial time points to estimate the bacterial load is recommended. (ii) Agar growth plates must be very dry before spot plating or else the individual 5 µL spots will merge. Either allow your agar plates to dry at room temperature for ∼2 days before use or place in a biosafety cabinet with the lids ajar for 2-4 hours before use. (iii) Select the lowest dilution with 10-25 discrete colonies to calculate the CFU of the wound. The same dilution does not need to be selected for each wound, as bacterial loads may be very different between treatment groups. (iv) We recommend plating the samples on both non-selective (e.g., LB agar) and selective and/or differential agar plates to assess both the colonization of the strain used to inoculate the wound, as well as any bacteria that may contaminate the wound during dressing changes.
7. **Study Termination**
  7.1 Anesthetize the pig and perform final wound imaging and surface bacteria sample collection as described in step 5.
  7.2 Perform euthanasia by injecting intravenous pentobarbital (100-120 mg/kg) and confirm expiration with the veterinarian before proceeding.
  7.3 Make a deep circular incision with a surgical scalpel around the wound boundary.
  7.4 Using Adson forceps to lift the wound edge and deepen the incision through the subcutaneous fat layer using Metzenbaum dissection scissors vertically until encountering the deep fascia (2-3 cm deep). Once at the deep fascia level, turn the direction of the scissors to start dissecting underneath the wound. There should be a clear separation of planes between the adipose tissue and the deep fascia layer.
  7.5 Place the excised tissue on a cutting board, keeping the wound orientation the same for all wounds. Cut the wound into strips (1-2 cm wide) using dissection scissors.
  7.6 Pin the central tissue strip on a 2×9 cm cork board and place in a 50 mL conical tube filled with HistoChoice® fixative.
  7.7 Place additional tissue strips into screw cap tubes and freeze for further biochemical/molecular analyses.
  7.8 Dispose of the animal corpse as a biohazard per institutional guidelines.
  **NOTE**: (i) Changes in skin appearance and temperature immediately after euthanasia can impact the evaluation of wound appearance, measurement of granulation tissue, and viability of bacteria collected from the wound surface. Therefore, these measurements should be performed on a live, anesthetized animal for consistent results. (ii) Ensure the entire tissue piece is immersed in HistoChoice® fixative. Due to tissue thickness, replace HistoChoice® 24 hours later and immerse for an additional 24 h to achieve optimum fixation.

**Figure 2:**
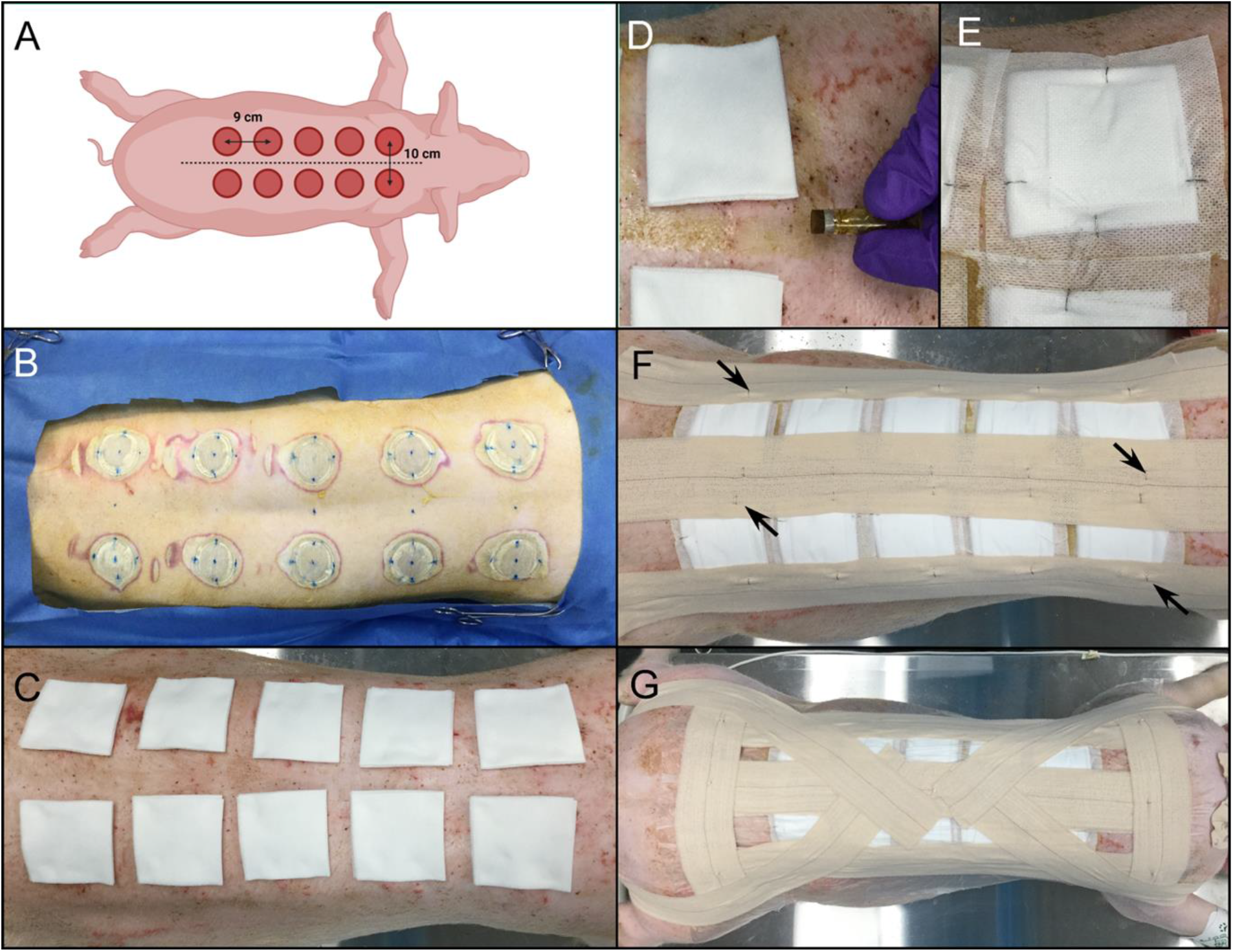
Wound placement design and dressing steps. **(A)** Diagram of wound placement. Pig is measured from the scapulae to the iliac crests (dotted lines) to determine optimal number and spacing of wounds paravertebrally, 5 cm from midline. **(B)** Appearance of skin post-burn creation. Note the blue dots marking the spine and wound placement to ensure even spacing of the wounds. **(C)** Wounds covered with 2×2” non-stick gauze pads, with fold towards midline. **(D)** Application of tincture of benzoin around the edges of the gauze pads. **(E)** Telfa dressing and gauze pad secured with skin staples. The staples should oppose each other and be perpendicular to the gauze edge. **(F)** Application of Elastikon® strips along the spine and on each side of the torso to reinforce the Telfa bandages. Staples are applied along the strips to anchor the strips to the dressings (black arrows). **(G)** Steri-drape dressings cover the prior dressing layers and are secured by Elastikon® strips along the spine, lateral to the dressings, and perpendicularly across the shoulders and hips. Staples are added to the perpendicular strips and down the central strip. An additional Elastikon® strip is placed from the center of the back, across the shoulder, under the neck, and across the other shoulder. Two final Elastikon® strips are applied from the center of the back and over the hip and butt.

**Figure 3:**
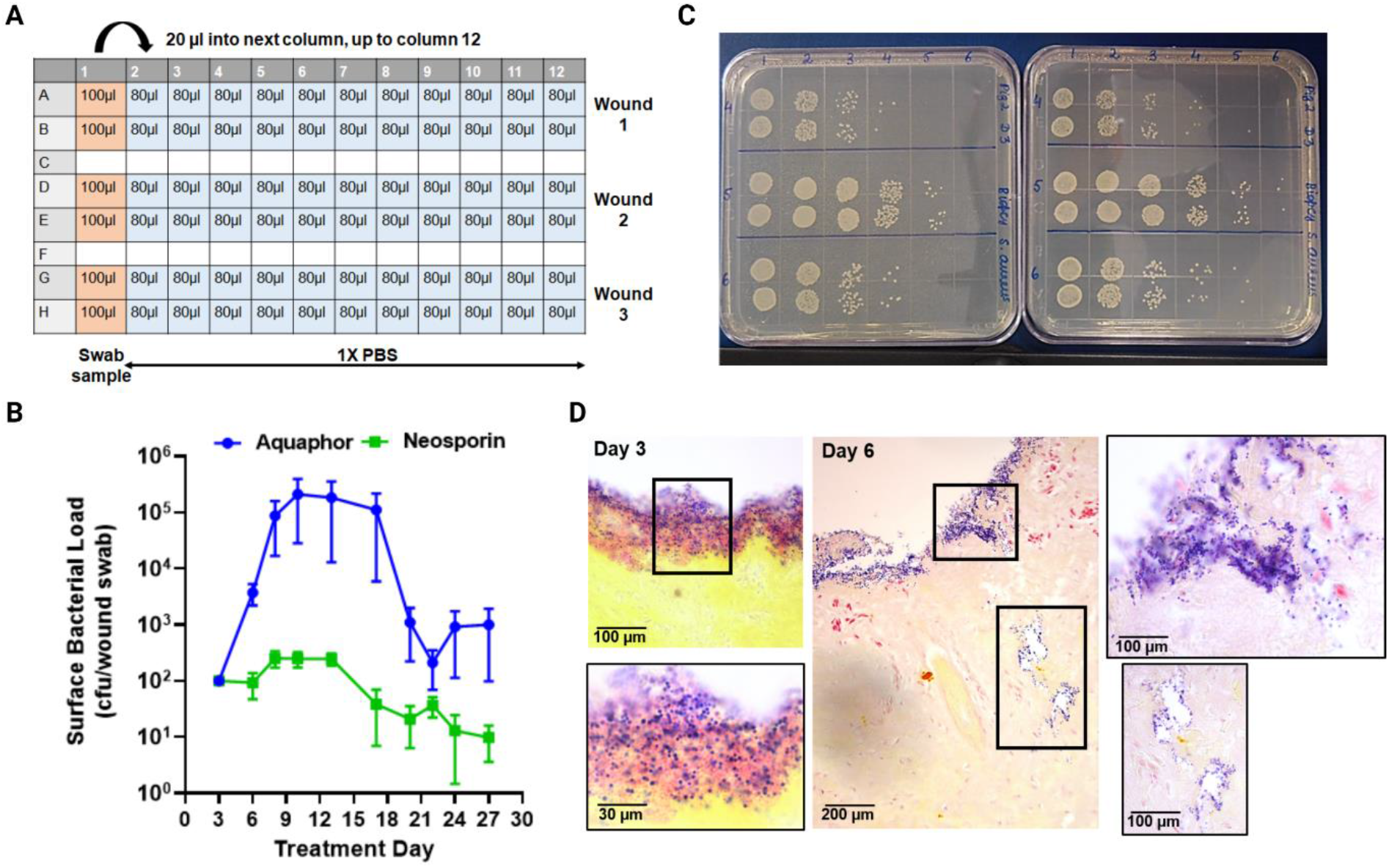
Assessment of bacterial colonization in burn wounds. **(A-C)** Culture assessment of surface bacterial colonization of burn wounds. **(A)** Dilution plate overview. An undiluted sample is aliquoted into column 1 of a sterile 96 well plate (100 µL, orange) in duplicate (e.g. rows A&B = wound 1, D&E = wound 2, G&H = wound 3). Sterile PBS is added to the remaining wells in each sample row (80 µL, blue). Serial dilutions are made by transferring 20 µL sample to the neighboring well to the right (e.g., column 1 to column 2, then column 2 to column 3, etc.). **(B)** Appearance of bacterial agar plates for dilutions 1-6 after spot plating and overnight growth. Using a multi-channel pipette, 5 µL from each well shown in (A) were spotted onto duplicate square agar plates and incubated overnight. **(C)** Quantification of the average viable CFU/wound determined by culture assessment over the duration of the experiment (n=36 wounds; mean±SEM). **(D)** Visualization of bacterial colonization by modified Gram stain of biopsy tissue samples from Day 3 and Day 6 post-wounding and inoculation. Insets show a higher magnification to highlight the surface colonization on Day 3 and invasive colonization on Day 6.

## ANTICIPATED RESULTS

### Bacterial colonization

Collection of surface swab samples and biopsies during dressing change procedures provides valuable information about the bacterial load on wound surfaces and in wound beds. In this study, treatment commenced three days after inoculation of wounds to promote the formation of a bacterial biofilm on the wound. Recovery of viable bacteria from the wound surface increased sharply between D3 to D6, and then precipitously dropped between D13 through D22 (**Fig. 3B**). We tested the efficacy of Neosporin, a known antibiotic ointment, against control (Aquaphor) in accelerating bacterial clearance from wounds. Results showed that Neosporin effectively cleared bacteria from wounds starting at D6 and the antibiotic treatment maintained an overall low surface bacterial load throughout the period of 27 days. *S. aureus* in tissue samples was visualized by Gram staining of fixed, paraffin-embedded tissue sections, as the dye stains the peptidoglycan-rich cell wall of the bacteria a dark purple (**Fig. 3D**). Increased surface bacterial load between D3 and D6 correlates with Gram stain visualization of *S. aureus* initially on the wound surface on D3, followed by localization of *S. aureus* under the wound eschar at later time points (**Fig. 3D**).

### Wound healing kinetics

Tissue biopsy samples collected from each wound once a week for histopathology and photographs of the wounds acquired at each dressing change over a period of 27 days provide a snapshot of the wound healing kinetics in the porcine model. H&E stains were analyzed and scored using a dermis score for assessment of tissue inflammation. (**Supplementary Table 1**). The dermis score constitutes grading features such as the presence of inflammatory cells, adipocytes, fibroblasts, hair follicles, and collagen fibers on a scale of 0 to 7 in tissue sections stained with H&E [12]. At earlier time points in this study, abundant adipocytes (day 3) and neutrophils (day 10) were observed in the dermis, indicative of an inflammatory phase. However, over time as the wounds healed in the last week of the study, tissues sections revealed the presence of fibroblasts and collagen deposition, indicating that the wound healing progressed to a tissue remodeling phase (**Fig. 4A-D**). Photographs acquired during the study were utilized to quantify the wound area compared to baseline (day 0 wounds). Wound area gradually increased until day 10, followed by a rapid decrease until day 22. The percentage of wound closure can also be quantified by subtracting the area of granulation tissue from the original wound area; however, this can only be performed after wound debridement (D17), as the initial eschar masks early granulation tissue (**Supplementary Fig. 3**).

**Figure 4:**
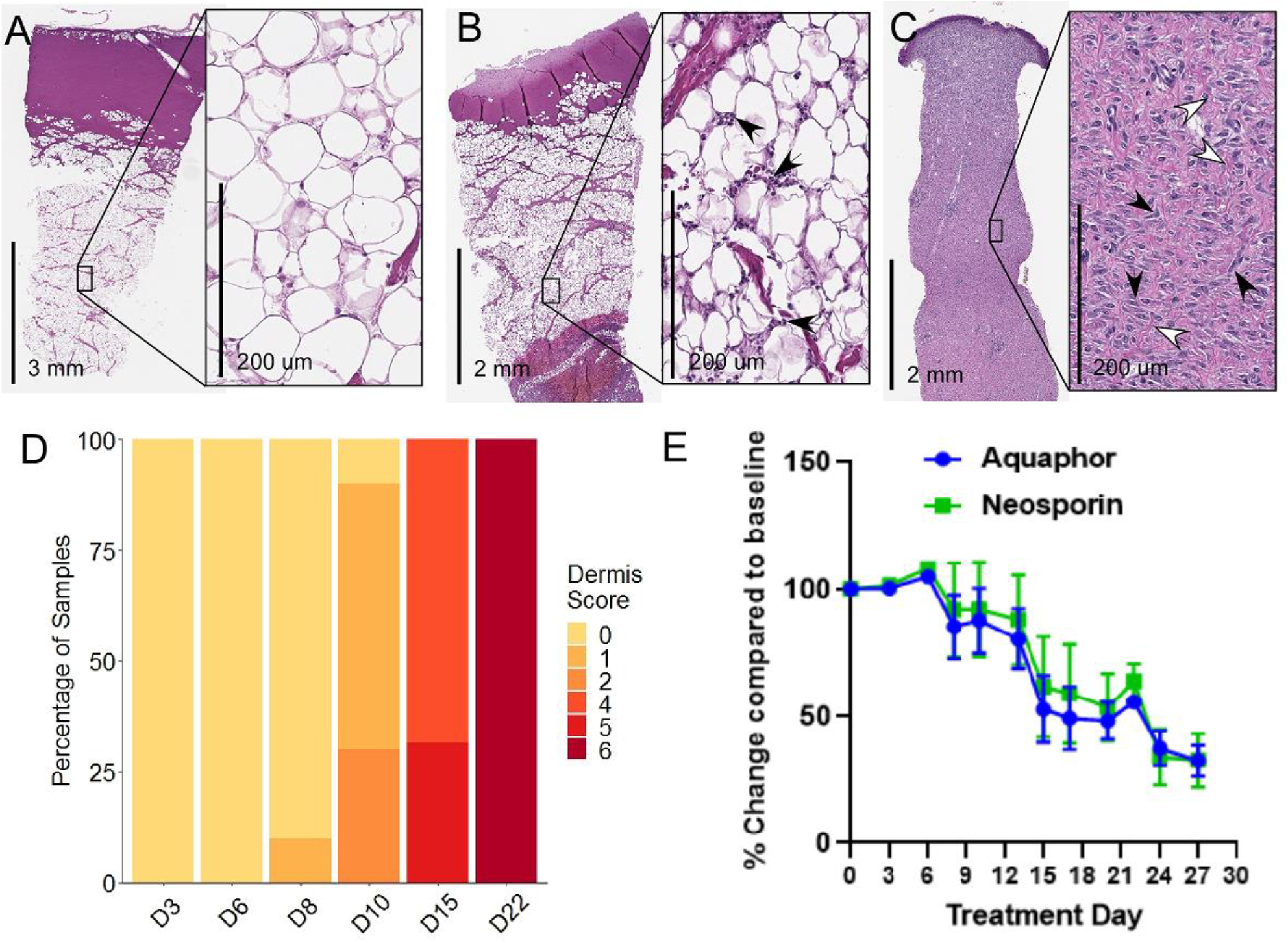
Wound healing assessment by wound closure and histologic scoring. **(A-C)** H&E stained wound biopsy tissue illustrating the progression of dermal regeneration after burn. **(A)** D3 biopsy showing an abundance of adipocytes in the dermal layer with minimal to no inflammatory cell infiltration. **(B)** D10 biopsy showing moderate inflammatory cell infiltration in the dermal layer, with numerous polymorphic neutrophils (black arrows). **(C)** D22 biopsy showing the disappearance of the dermal adipocytes, abundance of fibroblasts (black arrows) and collagen deposition (white arrows). **(D)** Dermis score progression (n=20 wounds) [13]. **(E)** Average wound area change over time compared to individual wound baseline measurements (n=36 wounds; mean±SEM).

## TROUBLESHOOTING

Common issues that could arise and possible solutions to consider are listed in **Table 1**.

**Table 1:**
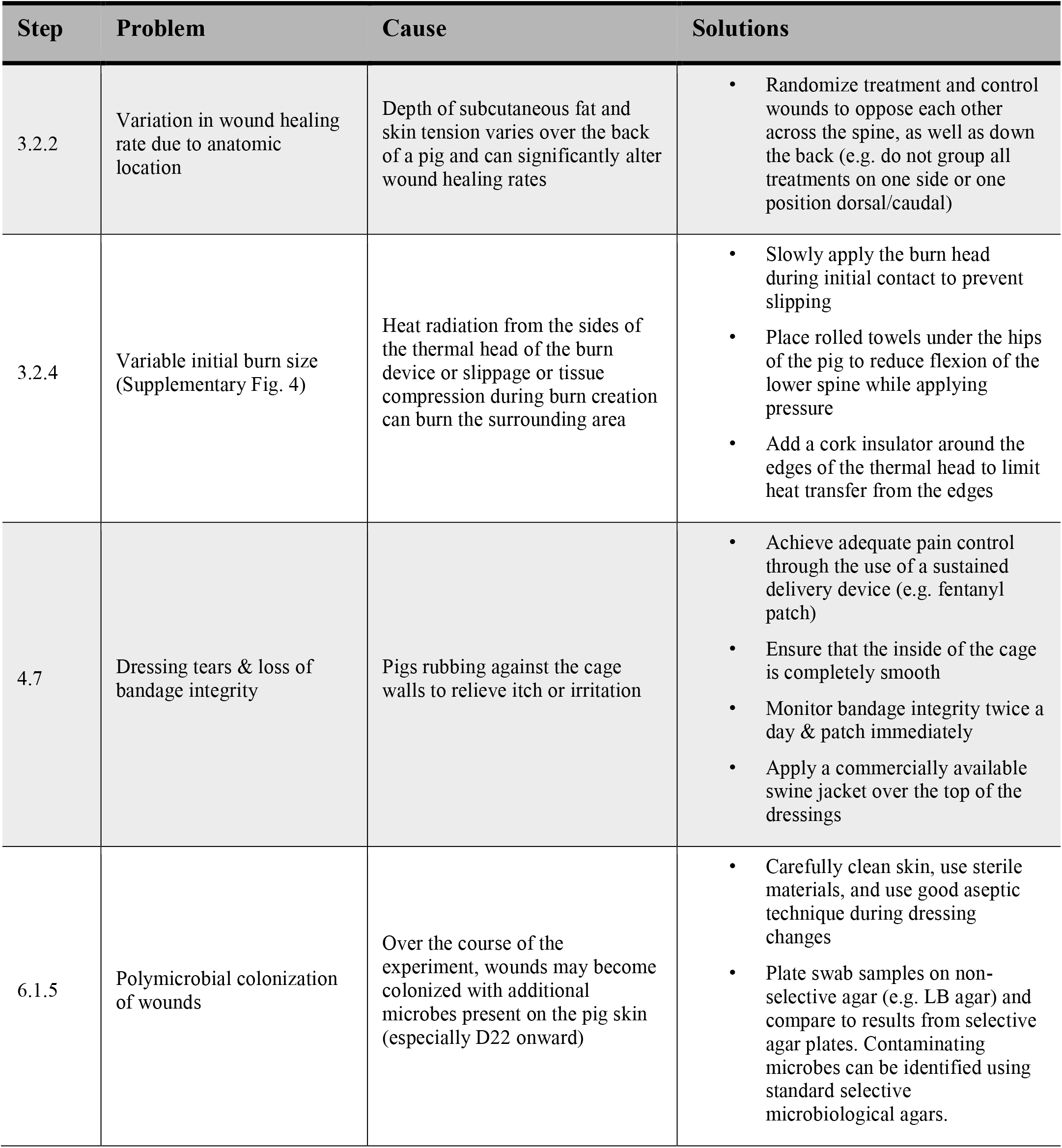
Commonly encountered problems, causes, and potential solutions.

1. **Experimental design and wound number estimation**.
  1.1 The maximum number of wounds per pig is primarily guided by the total available surface area, burn depth, wound size, and inter-wound distance required for bandaging each wound separately to reduce cross-contamination.
  1.2 The available surface area of the specific pig is determined by measuring the length of the spine from the scapula to iliac crest (**Fig. 2A**).
  1.3 Circular wounds 4 cm in diameter are applied symmetrically along the length of the spine, 5 cm from the midline, and a minimum of 5 cm between wound edges on the rostral/caudal axis.
  1.4 Once the wounding plan is designed, a plastic template can be created to facilitate the even spacing of the wounds (**Supplementary Fig. 2**).
  1.5 A Yorkshire pig 45-50 kg will have an available length of ∼45 cm, permitting 10 circular burn wounds 4 cm in diameter spaced 5 cm apart and 5 cm from the spinal cord.
2. **Tissue biopsy collection considerations**.
  2.1 The frequency of tissue sample collection needs to be balanced with the consideration that the accompanying injury may alter wound healing kinetics.
  2.2 We mitigated the impact of biopsy wounding by collecting samples once a week and from a similar anatomical location in each wound. Alternately, some studies dedicate wounds for collecting tissue biopsies and do not include these wounds in healing rate measurements.
  2.3 Biopsy collection becomes increasingly difficult in the later stages of wound healing and granulation and may result in bleeding during collection. Keep sterile small scissors and fine-tipped forceps ready to grasp the biopsy and separate the sample from the wound bed. Also have sterile gauze ready to place over the biopsy wound and apply firm pressure on the gauze with a finger to stop bleeding.
  2.4 Disinfect the forceps and scissors in between wounds to reduce cross-contamination.
  2.5 Bleeding during debridement can be controlled by applying firm pressure on the wound using a sterile gauze pad for a few minutes.

### Biosafety Considerations

All animal experiments and infectious agent handling must be performed in accordance with institutional and governmental ethics guidelines and regulations. This protocol was approved by the Institutional Animal Care and Use Committee and the Institutional Biosafety Committee of the Cleveland Clinic. A risk assessment must be performed in consultation with the biosafety and veterinary staff to determine the specific work practices to ensure the safety of personnel and animals within the facility. A resource for recommended biosafety practices is found in Biosafety in Microbiological and Biomedical Laboratories, 6^th^ edition [13].

## Abbreviations used

BSL2: Biosafety Level 2;
CFU: Colony Forming Unit;
H&E: Hematoxylin and Eosin;
OD: Optical Density;
PBS: Phosphate Buffered Saline;
PPE: Personal Protective Equipment;
TSB: Tryptic Soy Broth;
LB: Luria Broth;
MSA: Mannitol Salt Agar

## Acknowledgements

Such laborious experiments are only possible with the dedicated work of animal husbandry personnel and the veterinary technicians. We appreciate the tireless effort provided by our veterinary technicians Mary Lachowski, Jackie Kattar, and Orazio Terrano, as well as our husbandry technician Margaret Hanuschak. This research was supported by the Office of the Assistant Secretary of Defense for Health Affairs through the Congressionally Directed Medical Research Programs Peer Reviewed Medical Research Program under Award No. W81XWH1610439 (PR150299) and philanthropic support from the McDonald Family Trust. Opinions, interpretations, conclusions, and recommendations are those of the authors and are not necessarily endorsed by the funders. The graphical abstract, Figure 1 and the cartoon in Figure 2A was created with BioRender.com.

## References

1. Jeschke M, van Baar M, Choudhry M, Chung K, Gibran N, Logsetty S. Burn injury. Nature Reviews Disease Primers. 2020;6(11). doi: https://doi.org/10.1038/s41572-020-0145-5. PubMed Central PMCID: PMC32054846.

2. Seaton M, Hocking A, Gibran NS. Porcine models of cutaneous wound healing. ILAR J. 2015;56(1):127–38. doi: 10.1093/ilar/ilv016. PubMed PMID: 25991704.

3. Moins-Teisserenc H, Cordeiro DJ, Audigier V, Ressaire Q, Benyamina M, Lambert J, et al. Severe Altered Immune Status After Burn Injury Is Associated With Bacterial Infection and Septic Shock. Front Immunol. 2021;12:586195. doi: 10.3389/fimmu.2021.586195. PubMed PMID: 33737924; PubMed Central PMCID: PMCPMC7960913.

4. Church D, Elsayed S, Reid O, Winston B, Lindsay R. Burn wound infections. Clin Microbiol Rev. 2006;19(2):403–34. doi: 10.1128/CMR.19.2.403-434.2006. PubMed PMID: 16614255; PubMed Central PMCID: PMCPMC1471990.

5. Lachiewicz AM, Hauck CG, Weber DJ, Cairns BA, van Duin D. Bacterial Infections After Burn Injuries: Impact of Multidrug Resistance. Clin Infect Dis. 2017;65(12):2130–6. doi: 10.1093/cid/cix682. PubMed PMID: 29194526; PubMed Central PMCID: PMCPMC5850038.

6. Summerfield A, Meurens F, Ricklin ME. The immunology of the porcine skin and its value as a model for human skin. Mol Immunol. 2015;66(1):14–21. doi: 10.1016/j.molimm.2014.10.023. PubMed PMID: 25466611.

7. Dawson HD, Loveland JE, Pascal G, Gilbert JG, Uenishi H, Mann KM, et al. Structural and functional annotation of the porcine immunome. BMC Genomics. 2013;14:332. doi: 10.1186/1471-2164-14-332. PubMed PMID: 23676093; PubMed Central PMCID: PMCPMC3658956.

8. Singh M, Nuutila K, Minasian R, Kruse C, Eriksson E. Development of a precise experimental burn model. Burns. 2016;42(7):1507-12. doi: 10.1016/j.burns.2016.02.019. PubMed PMID: 27450518.

9. Wang XQ, Kempf M, Liu PY, Cuttle L, Chang HE, Kravchuk O, et al. Conservative surgical debridement as a burn treatment: supporting evidence from a porcine burn model. Wound Repair Regen. 2008;16(6):774–83. doi: 10.1111/j.1524-475X.2008.00428.x. PubMed PMID: 19128248.

10. Cuttle L, Kempf M, Phillips GE, Mill J, Hayes MT, Fraser JF, et al. A porcine deep dermal partial thickness burn model with hypertrophic scarring. Burns. 2006;32(7):806–20. doi: 10.1016/j.burns.2006.02.023. PubMed PMID: 16884856.

11. Gusarov I, Shatalin K, Starodubtseva M, Nudler E. Endogenous nitric oxide protects bacteria against a wide spectrum of antibiotics. Science. 2009;325(5946):1380–4. doi: 10.1126/science.1175439. PubMed PMID: 19745150; PubMed Central PMCID: PMCPMC2929644.

12. Guo HF, Mohd Ali R, Abd Hamid R, Chang SK, Zainal Z, Khaza’ai H. A new histological score grade for deep partial-thickness burn wound healing process. Int J Burns Trauma. 2020;10(5):218-24. PubMed PMID: 33224609; PubMed Central PMCID: PMCPMC7675209.

13. Centers for Disease Control and Prevention, National Institutes of Health. Biosafety in Microbiological and Biomedical Laboratories. 6^th^ Edition, June 2020.

