## Supplemental Table & Figures for "Development of a Reproducible Porcine Model of Infected Burn Wounds"

Supplementary Table 1
Supplementary Figures 1-4

**Supplementary Table 1. Histological dermal score system after burn injury** [12].

| Dermis Score | Adipocytes | Inflammatory Cells | Fibroblast | Collagen Deposition | Hair Follicles |
|---|---|---|---|---|---|
| **0** | +++ | + | – | – | – |
| **1** | ++ | ++ | + | – | – |
| **2** | + | ++ | + | – | – |
| **3** | + | +++ | + | – | – |
| **4** | + | +++ | ++ | + | – |
| **5** | – | ++ | +++ | ++ | – |
| **6** | – | + | +++ | ++ | – |
| **7** | – | + | ++ | +++ | + |

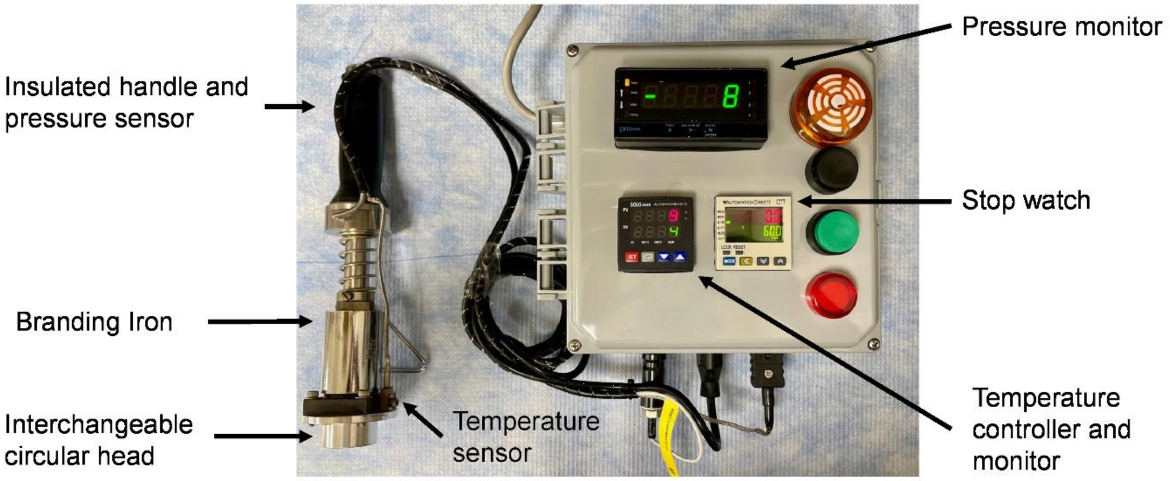


**Supplementary Figure 1: Custom burn wound device.** Features of the device that regulate delivery of thermal energy and pressure to create consistent burn wounds. Plans are available upon request.

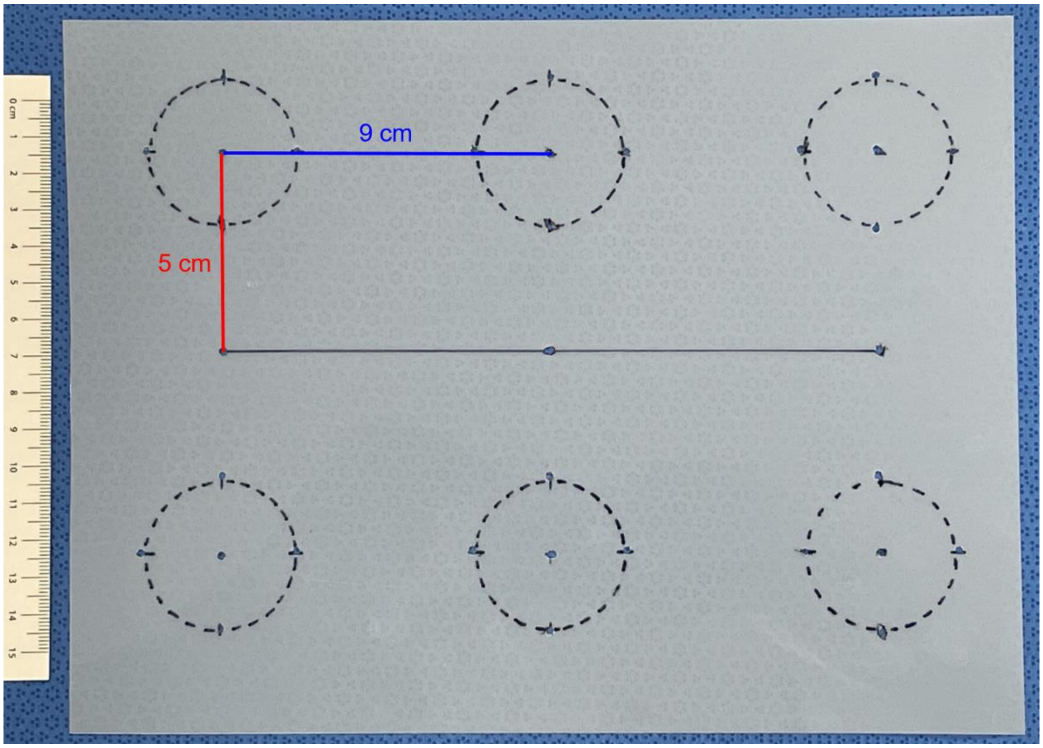


**Supplementary Figure 2: Wound creation template.** A plastic stencil to mark consistent spacing of wound areas from the spine (center line) and each other longitudinally. The template is perforated to permit the marking of the pig skin with a skin marker/Sharpie.

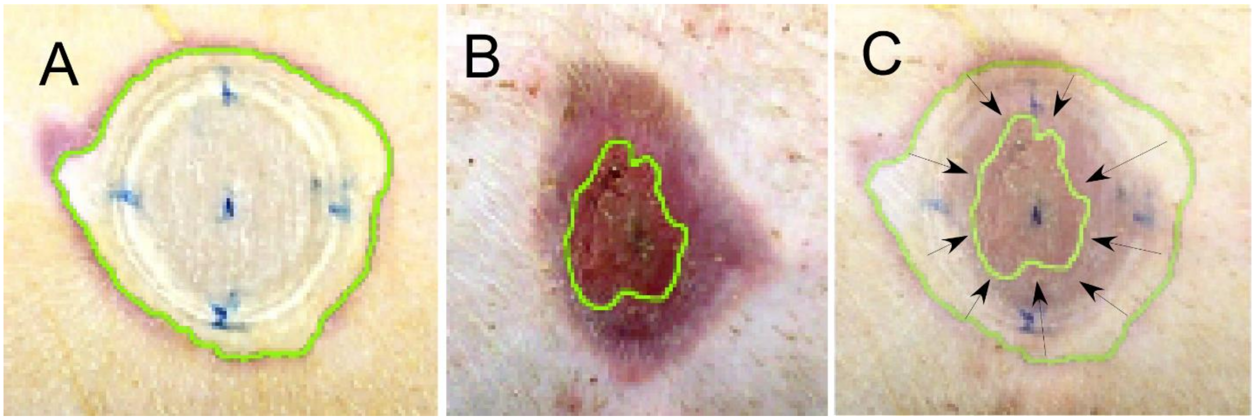


**Supplementary Figure 3: Wound closure measurement. (A)** Baseline wound area measured on D0 (green outline). **(B)** Granulation area (green outline) measured on D27. **(C)** Percent of wound closure calculated by subtracting the granulation area from the baseline wound area, then dividing by the baseline wound area, and multiplying by 100.

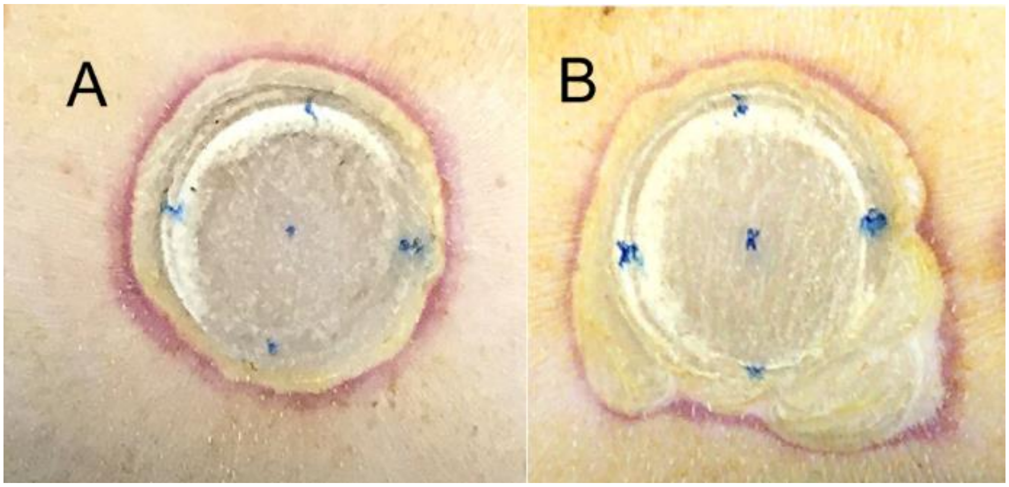


**Supplementary Figure 4: Variation in burn wound creation. (A)** Uniform burn with minimal heat radiation to the surrounding tissue. **(B)** Heat radiation damaging the surrounding tissue resulting in a different baseline wound area.
